# *Staphylococcus aureus* infects osteoclasts and replicates intracellularly

**DOI:** 10.1101/638528

**Authors:** Jennifer L Krauss, Philip M Roper, Anna Ballard, Chien-Cheng Shih, James AJ Fitzpatrick, James E Cassat, Pei Ying Ng, Nathan J Pavlos, Deborah J Veis

## Abstract

Osteomyelitis (OM), or inflammation of bone tissue, occurs most frequently as a result of bacterial infection and severely perturbs bone structure. The majority of OM is caused by *Staphylococcus aureus*, and even with proper treatment, OM has a high rate of recurrence and chronicity. While *S. aureus* has been shown to infect osteoblasts, persist intracellularly, and promote the release of pro-osteoclastogenic cytokines, it remains unclear whether osteoclasts (OCs) are also a target of intracellular infection. In this study, we examined the interaction between *S. aureus* and OCs, demonstrating internalization of GFP-labeled bacteria by confocal microscopy, both *in vitro* and *in vivo*. Utilizing an intracellular survival assay and flow cytometry during OC differentiation from bone marrow macrophages (BMMs), we found that the intracellular burden of *S. aureus* increases after initial infection in cells with at least 2 days of exposure to the osteoclastogenic cytokine receptor activator of nuclear factor kappa-B ligand (RANKL). Presence of dividing bacteria was confirmed via visualization by transmission electron microscopy. In contrast, undifferentiated BMMs, or those treated with interferon-γ or IL-4, had fewer internal bacteria, or no change, respectively, at 18 hours post infection, compared to 1.5 hours post infection. To further explore the signals downstream of RANKL, we manipulated NFATc1 and alternative NF-κB, which controls NFATc1 and other factors affecting OC function, finding that intracellular bacterial growth correlates with NFATc1 levels in RANKL-treated cells. Confocal microscopy in mature OCs showed a range of intracellular infection that correlated inversely with *S. aureus* and phagolysosome colocalization. The ability of OCs to become infected, paired with their diminished bactericidal capacity compared to BMMs, could promote OM progression by allowing *S. aureus* to evade initial immune regulation and proliferate at the periphery of lesions where OCs and bone remodeling are most abundant.

**Author Summary:** The inflammation of bone tissue is called osteomyelitis, and most cases are caused by an infection with the bacterium *Staphylococcus aureus*. To date, the bone building cells, osteoblasts, have been implicated in the progression of these infections, but not much is known about how the bone resorbing cells, osteoclasts, participate. In this study, we show that *S. aureus* can infect osteoclasts and proliferate inside these cells, whereas macrophages, immune cells related to osteoclasts, destroy the bacteria. These findings elucidate a unique role for osteoclasts to harbor bacteria during infection, providing a possible mechanism by which bacteria could evade destruction by the immune system. Therapeutic interventions that target osteoclasts specifically might reduce the severity of OM or improve antibiotic responses.

## Introduction

Although the name technically refers to the inflammation of the marrow cavity, the term osteomyelitis (OM) most frequently is used to indicate infection of the bone itself. At the center of infectious OM lesions, bone is frequently lost by necrosis, forming a devascularized segment of bone known as a sequestrum. Osteoclasts (OCs) are recruited to the site via inflammatory cytokine release and expand the area of bone loss. New bone is formed on the periosteum in the body’s attempt to isolate the infection, generating an involucrum. Thus, the normal balance between bone formation and resorption is disrupted in OM, leading to pathologic fractures and deformities [1]. Compared to other tissues, bone infections are especially damaging and intractable, with treatment often involving prolonged antibiotics paired with surgical debridement [2]. However, even with proper treatment, OM has a high recurrence rate, leading to a chronic debilitating condition [3, 4]. OM is predominantly divided into two broad categories: 1) acute hematogenous OM caused by bacteria seeding directly into bone from the circulation, and 2) secondary OM originating from a contiguous source like a soft tissue infection or orthopedic implant, or following open fracture [1, 5]. Acute hematogenous OM is most common in children, with 85% of cases occurring in children under 17 years of age, whereas secondary OM infections are more common in older adults, especially those undergoing orthopedic surgery [5, 6]. Regardless of the type of OM, most cases are caused by *Staphylococcus aureus* [6, 7].

Despite the clinical significance of OM infections, there remains a dearth of knowledge as to the mechanisms underlying the etiopathology of the disease. To this point, most of the basic science work on OM has focused on characterizing changes to the structure of the bone [8, 9, 10], elucidating bacterial survival strategies [11, 12], or examining the role of osteoblasts in promoting infection [11, 13, 14, 15]. *S. aureus* has the ability to infect osteoblasts, persist intracellularly, and induce the release of osteoclastogenic and inflammatory cytokines, leading to OC recruitment and differentiation at the site of infection [9, 13, 14, 16]. Investigation into the role of OCs in OM has focused mainly on determining the effects of *S. aureus* infection on bone resorption, while neglecting to examine whether OCs could be the target of infection, as well. [17, 18, 19, 20].

OCs are differentiated from the myeloid lineage in a process that involves RANKL signaling to alternative NF-κB signaling through NIK and RelB, activating NFATc1 [21, 22, 23]. However, despite their shared lineage, OCs show a decreased release of inflammatory cytokines and nitric oxide when challenged with bacteria as compared to macrophages [19]. Macrophages attempt to destroy internalized *S. aureus* via phagolysosomal acidification [24], but *S. aureus* can also persist within macrophages through intraphagolysosomal replication and escape [25, 26]. Yet, the ability of OCs to control *S. aureus* infection is not known. In this study, we show that *S. aureus* is not only able to infect OCs, but also proliferates within them, avoiding lysosomal compartments. These results highlight osteoclasts’ potential facilitation of *S. aureus* immune evasion, which could be an important mechanism in the propagation of OM infections, as well as explain the propensity for OM recurrence. This study illuminates a novel function of osteoclasts in OM as an intracellular reservoir allowing bacterial proliferation, in addition to their ability to modulate bone structure through perturbed remodeling.

## Results

### *S. aureus* resides within osteoclasts *in vivo* and *in vitro*

Although previous studies have shown that *S. aureus* invasion of osteoblasts leads to RANKL expression and robust OC recruitment at sites of bone infection [13, 14], it remains unclear whether OCs are cellular targets for intracellular infection. To demonstrate the intracellular presence of *S. aureus* within OCs *in vivo*, we injected GFP expressing-*S. aureus* (USA300-lineage strain LAC) subcutaneously over the periosteum of calvaria in TRAP-tdTomato (herein referred as TRAP^Red^) reporter mice [27]. Prior to microbial challenge, RANKL was injected daily over the calvaria for 5 days to recruit OCs and their precursors to the subsequent site of infection. At 24 hours post-infection (hpi), calvaria were harvested and histological sections were examined by confocal microscopy to visualize bacteria residing within OCs, since both are fluorescently labeled. As shown in Figure 1A, GFP-expressing *S. aureus* can clearly be found localized within TRAP^Red^ OCs. Murine OCs were also generated on bone slices *in vitro*, then infected with GFP-labeled *S. aureus* and imaged 18 hours later. Again, bacteria were found within OCs (Fig 1B).

**Fig 1.**
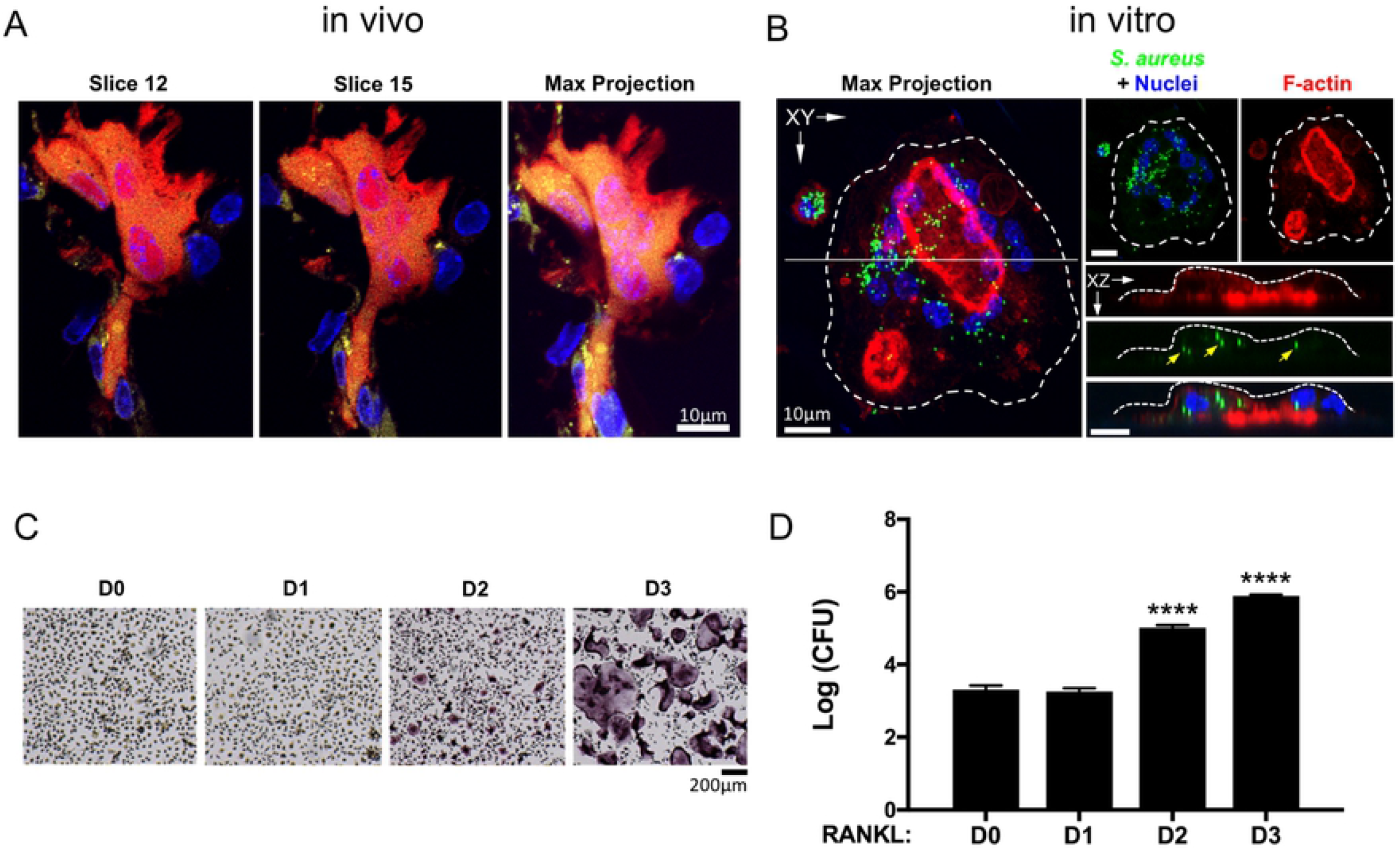
*S. aureus* resides within OCs *in vivo* and *in vitro*. (A) Confocal microscopy of histological sections reveals internalized GFP+ *S. aureus* inside TRAP^Red^ OCs within mouse calvarium (yellow) and (B) within OCs differentiated from BMMs with three days of RANKL and M-CSF on glass coverslips (green puncta). (C) TRAP staining of BMMs differentiated towards OCs for up to three days showing multiple TRAP+ mononuclear cells at D2 and numerous TRAP+ multinuclear fully differentiated OCs at D3. (D) Enumerated colony forming units (CFUs) grown on tryptic soy agar from lysates of BMMs differentiated towards OCs for up to three days and infected with *S. aureus* for 18 hours, with gentamicin killing of extracellular bacteria. ****p<0.0001 by one-way ANOVA with Tukey’s post-hoc test, compared to D0. n=3 technical replicates, representative > 5 biological replicates.

In order to demonstrate whether the observed intracellular bacteria were viable, we used a gentamicin-based protection assay. Murine bone marrow macrophages (BMMs) were differentiated with RANKL for up to 3 days (Fig 1C) and infected with *S. aureus* for 30 minutes at a multiplicity of infection (MOI) of 1:1, after which extracellular bacteria were killed by the addition of gentamicin for 1 hour. Infected cells were lysed following 16.5 hours of additional culture (at 18 hpi), and colony forming units (CFUs) enumerated. While exposure of BMMs to RANKL for 1 day had no effect on the number of bacteria recovered, 2 days of differentiation in RANKL (lineage-committed TRAP+ preosteoclasts [preOCs]) led to ~100-fold increased bacteria load, and 3 days of RANKL (fully differentiated OCs) caused a ~500-fold change, compared to no RANKL (undifferentiated BMMs) (Fig 1D). In order to preclude any confounding effects that may be specific to gentamicin treatment in our assay, we repeated the 18 hpi CFU assay during OC differentiation with lysostaphin as the bactericidal agent instead of gentamicin. We found the patterns of increased intracellular bacterial load in OCs at 18 hpi were the same in our lysostaphin protection assay as with our gentamicin protection assay (S1 Fig).

### *S. aureus* proliferates within osteoclasts *in vitro*

BMMs have innate immune activity and have previously been shown to phagocytose and kill intracellular bacteria [24]. Using the gentamycin protection assay, we confirmed that our BMMs behave similarly, finding that the number of intracellular *S. aureus* at 18 hpi is ~10-fold lower than at 1.5 hpi, immediately after removal of extracellular bacteria by antibiotic treatment (Fig 2A, D0). In contrast, bacterial recovery from RANKL-treated cultures (D2 or D3) demonstrated a dramatic increase between 1.5 and 18 hpi, suggesting that *S. aureus* can proliferate within OCs. There was no difference in the ability of any of the cultures to internalize *S. aureus*, as the 1.5 hpi bacterial counts were similar across groups. In order to rule out potential differential effects of OCs grown on culture-treated plastic, we confirmed similar results in OCs grown and infected on bone chips *in vitro* (S2 Fig). Next, to establish whether these observations were merely an artifact of murine OCs, we infected primary human OCs generated from peripheral blood CD14^+^ monocytes. Consistent with the data in the mouse system, human OCs significantly promote intracellular replication of *S. aureus* (Fig 2B).

**Fig 2.**
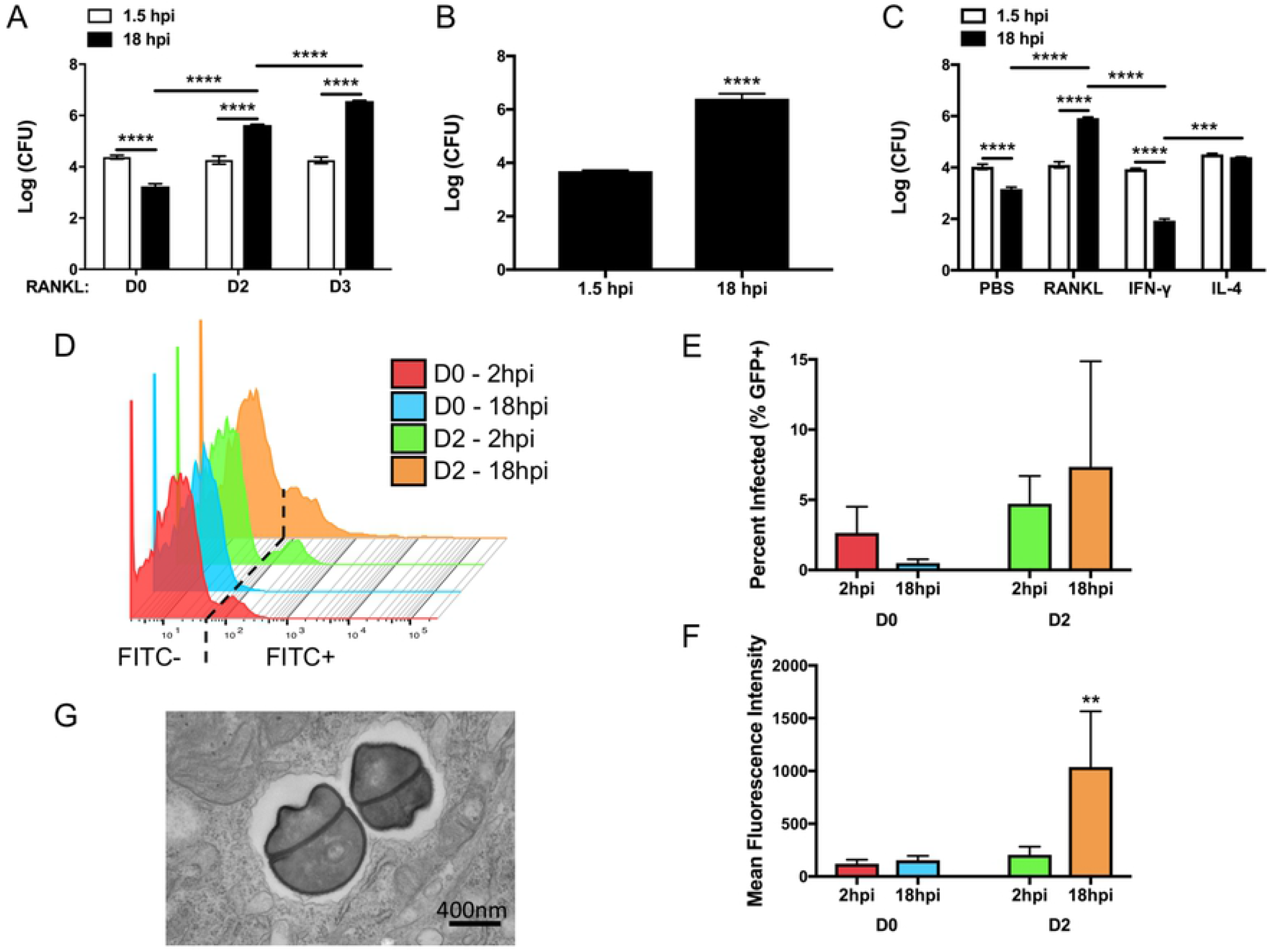
*S. aureus* proliferates within OCs *in vitro*. BMMs were treated with RANKL for up to three days and subjected to the gentamicin protection assay. (A) Colony forming units (CFU) from lysates of infected cells after OC differentiation for 0, 2, or 3 days. Cells were lysed immediately after gentamycin exposure (1.5 hpi) or after an additional 16.5 hour in osteoclastogenic (D2, D3) or control media (D0). n=3 technical replicates, representative > 5 biological replicates. (B) CFU from lysates of infected human CD14+ monocytes isolated from peripheral blood and differentiated into OCs for 3 days before infection. n=3 biological replicates. (C) CFU from lysates of infected BMMs after exposure for 2 days to either PBS, RANKL, IFN-γ or IL-4. n=3 technical replicates, representative of 3 biological replicates. (D) Offset histogram of flow cytometric data from infected cells differentiated in RANKL for 0 or 2 days, then infected with GFP+ *S. aureus* at an MOI of 1:1. Infected cells were detected by an increased signal in the FITC channel. The threshold for FITC+ is depicted by the dashed line, based on cells infected with GFP-bacteria. n=3 biological replicates. (E) Infected cells (GFP+) are shown as a percentage of total cells, as measured via flow cytometry. (F) Mean fluorescence intensity (MFI) as measured from the FITC+ fraction of cells via flow cytometry. (G) Transmission electron micrograph of dividing *S. aureus* in a membrane-bound compartment inside an OC. **p<0.01, ****p<0.0001 by two-way ANOVA with Tukey’s post-hoc test (A, C, F) or student’s t-test (B).

Since the antimicrobial properties of myeloid cells are altered by cytokine exposure, we sought to determine whether RANKL was unique in its ability to promote intracellular bacterial growth. To address this question, we polarized BMMs for 2 days with IFN-γ, towards an antimicrobial M1 phenotype, or with IL-4, towards an M2 phenotype expected to have reduced antimicrobial defense, and then compared these to RANKL-stimulated BMMs (i.e. preOC). In all cases, levels of bacteria were similar at 1.5 hpi, indicating consistent internalization (Fig 2C). IFN-γ treatment led to higher levels of killing than observed in control PBS-treated BMMs, as expected for M1-polarized cells. In the IL-4 stimulated, M2-polarized cells, we found that *S. aureus* are present at the same levels 1.5 and 18 hpi, indicating persistence rather than intracellular killing or proliferation. Thus, RANKL initiates a distinct program that alters the antimicrobial profile of BMMs as they differentiate toward OCs.

To further explore the fate of bacteria after internalization, we again utilized GFP-labeled *S. aureus*, this time for flow cytometry. BMMs or preOCs grown in RANKL for 2 days were infected at an MOI of 1:1, extracellular bacteria killed with gentamycin as before, and then cells were fixed at 2 or 18 hpi. At this low MOI, a minority of cells bear detectable levels of GFP+ bacteria at 2 hpi, and the level is similar between BMMs and preOCs (Fig 2D, 2E), as we observed in the bulk CFU assay (Fig 2A, 2C). By 18 hpi, the fraction of GFP+ BMMs decreased similar to previous observations, although it did not reach statistical significance. There was no statistically significant change in the percent of GFP+ D2 preOCs at 18 hpi, although there was higher variability between biological replicates. Very few bacteria were detected in the culture media at 12-18 hpi, suggesting it is unlikely that cell lysis and new infection of adjacent cells occurs at a high enough frequency to affect our readouts (S1 Table). We next plotted the mean fluorescence intensity (MFI) of the GFP+ populations in each culture (Fig 2F). There was no change in the BMMs with time, suggesting that the few cells that were unable to clear bacteria nevertheless restrained their growth. In contrast, the MFI of GFP+ preOCs increased ~4-fold between 2 and 18 hpi, indicating a greater number of bacteria per cell at the later timepoint. Further, the histogram (Fig 2D) shows a population of preOC at 18 hpi with a very high MFI (up 100-500-fold) compared to any other culture condition, suggesting that a subset of these cells allows rapid intracellular replication of *S. aureus*. Supporting our conclusion that the bacteria are expanding within RANKL-treated cells, transmission electron microscopy demonstrates dividing bacteria in mature OCs at 18 hpi (Fig 2G).

### Proliferative capacity of *S. aureus* within osteoclasts is NFATc1-dependent in response to RANKL

Having demonstrated that *S. aureus* proliferates robustly within differentiated OCs, we next wanted to determine whether deficiency of NFATc1, the master transcriptional regulator of OC formation, would modulate intracellular bacterial levels in response to RANKL. To this end, *Nfatc1^fl/fl^* mice were mated to an inducible *Mx1-cre* transgenic line and Poly I:C was used to conditionally delete *Nfatc1* one month prior to harvest of BMMs (Fig 3A). Consistent with the critical function of NFATc1 in driving OC differentiation (Fig 3B), we find that *S. aureus* was unable to replicate in NFATc1-deficient cells in response to RANKL stimulation (Fig 3C, D2 condition). In contrast, we found no difference in the microbial load between unstimulated Ctrl and NFATc1-deficient BMMs, a result consistent with the low basal expression of NFATc1 in the absence of RANKL (Fig 3A).

**Fig 3.**
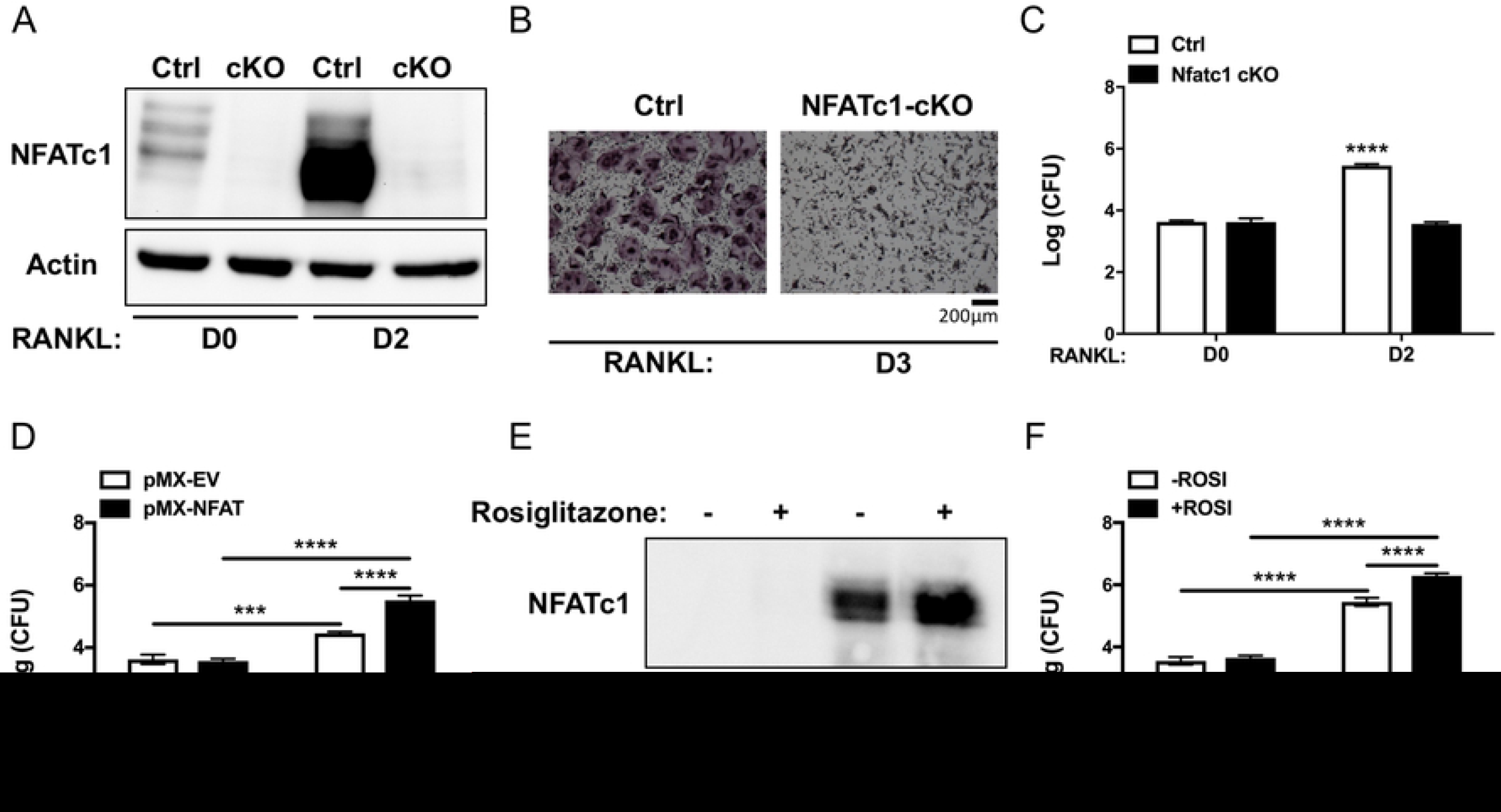
Intracellular growth of *S. aureus* is NFATc1-dependent in response to RANKL. (A) BMMs harvested from NFATc1 conditional knockout animals (cKO) show no basal (D0) or induced (D2) NFATc1 protein by western blot as compared to littermate control mice BMMs (Ctrl). (B) TRAP staining after 3 days of RANKL exposure demonstrates failure of NFATc1-cKO BMMs to form TRAP+ multinuclear OCs. (C) CFU from lysates of Ctrl or NFATc1 cKO BMMs differentiated into OCs for 0 or 2 days and subjected to the gentamicin protection assay. All bars represent 18 hours post infection (hpi). White bars, Ctrl; black bars, NFATc1 cKO cells. (D) CFU from lysates of wild-type (WT) BMMs transfected with empty vector (pMX-EV, white bars) or NFATc1 overexpressing vector (pMX-NFAT, black bars) at 18hpi, showing positive effect of NFATc1 at D2. (E) Rosiglitazone treatment of WT BMMs increases NFATc1 induction more than RANKL alone as measured by western blot of nuclear extracts. H3, histone 3 antibody. (F) CFU from lysates of WT BMMs treated with Rosiglitazone (+ROSI, black bars) or untreated (-ROSI, white bars) at 18hpi. n=3 biological replicates. ***p<0.001, ****p<0.0001 by two-way ANOVA with Tukey’s post-hoc test.

Because we found that RANKL-induced NFATc1 promotes intracellular replication of *S. aureus*, we evaluated whether forced expression of this molecule would further enhance microbial expansion. For this purpose, we cloned *Nfatc1* cDNA into the pMX retroviral vector and transduced Wt BMMs. Following blasticidin selection, transduced cells were cultured with M-CSF alone or under osteoclastogenic conditions and challenged with *S. aureus*. Using this gain-of-function approach, we demonstrate that ectopic expression of NFATc1, which accelerates OC differentiation, significantly increased microbial burden when compared to cells transduced with empty vector under osteoclastogenic conditions (Fig 3D). Surprisingly, overexpression of NFATc1 in BMMs, in the absence of RANKL signaling, failed to support replication, as differences in microbial load were comparable to empty vector control. Similarly, treatment with rosiglitazone, a peroxisome proliferator-activated receptor-γ agonist that augments RANKL-induced NFATc1 levels when combined with RANKL (Fig 3E), led to a significant increase in intracellular microbial accumulation relative to untreated OCs (Fig 3F). However, in the absence of RANKL, NFATc1 expression was not induced and intracellular microbial counts were comparable between rosiglitazone treated and untreated BMMs (Fig 3F). Thus, using both genetic and pharmacological approaches, intracellular proliferation of *S. aureus* within OCs is NFATc1-dependent in response to RANKL stimulation.

### Alternative NF-κB promotes intracellular expansion of *S. aureus*

Although our data demonstrate a clear role for NFATc1 in facilitating intracellular growth of *S. aureus* in OCs, we found that microbial burden was unaltered in NFATc1 overexpressing BMMs compared to empty vector controls, suggesting that other RANKL-induced signaling pathways are also essential for the effect. Previously, our group has demonstrated that the alternative NF-κB signaling pathway is highly upregulated by RANKL stimulation and promotes OC formation [22, 23, 28]. In order to investigate the importance of alternative NF-κB signaling in promoting *S. aureus* intracellular expansion, we utilized cells deficient in the alternative NF-κB central upstream kinase NIK or the downstream transcription factor RelB. Loss of either NIK or RelB significantly decreases the ability of *S. aureus* to replicate intracellularly (Fig 4A, 4B).

**Fig 4.**
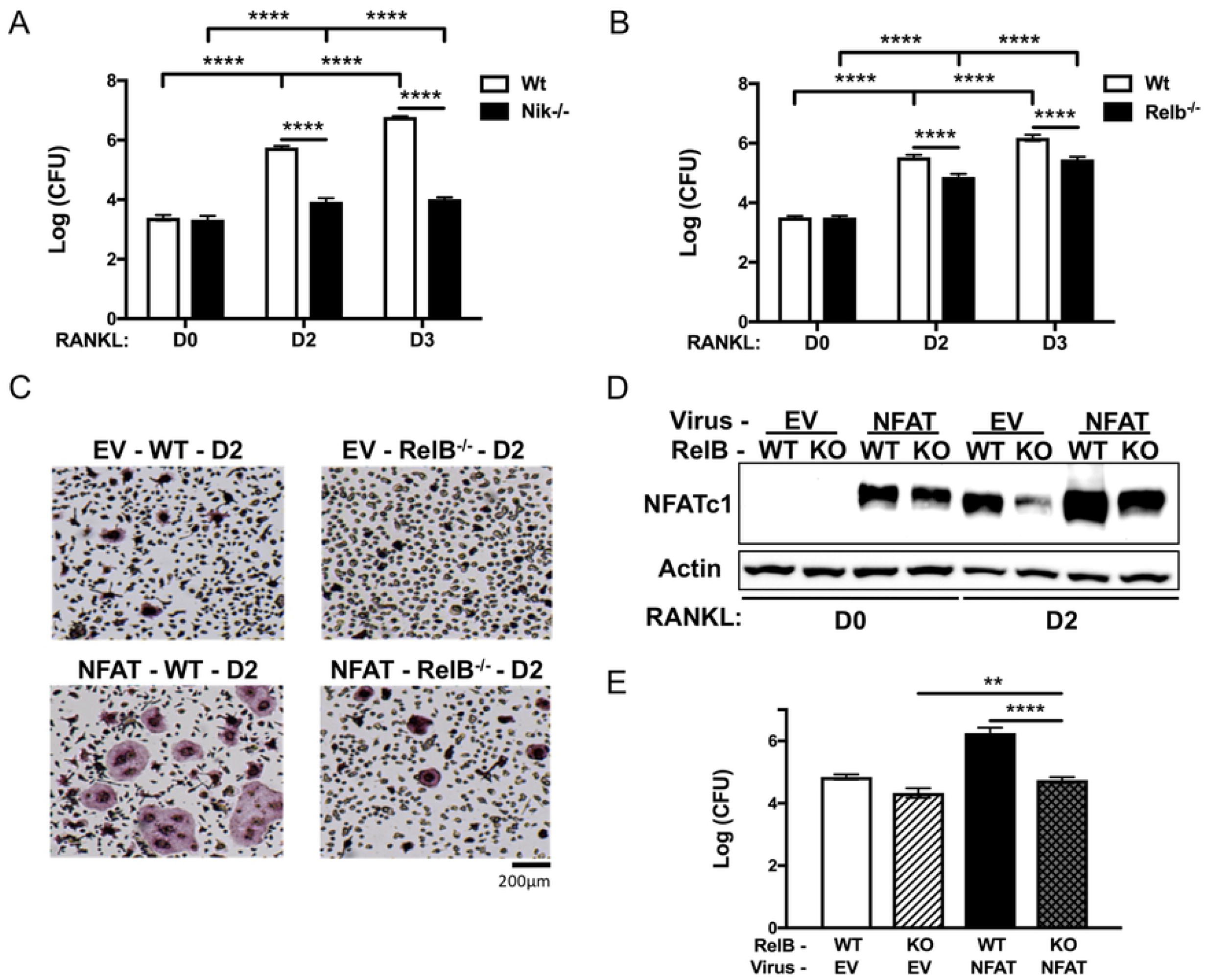
Alternative NF-κB in OCs promotes intracellular *S. aureus* replication *in vitro*. (A) BMMs harvested from NIK knockout mice (Nik−/−, black bars) and (B) RelB knockout mice (Relb−/−, black bars) were differentiated into OCs for 0, 2, or 3 days and subjected to the gentamicin protection assay. CFU of intracellular *S. aureus* proliferation were examined at 18 hpi. (C) TRAP stained images of WT or Relb−/− BMMs transduced with empty vector (EV) or NFATc1 overexpressing virus (NFAT) and differentiated for 48 hours. (D) Western blot of NFATc1 protein expression from WT or Relb−/− (KO) BMMs transduced with EV or NFAT and differentiated with RANKL for 0 (D0) or 2 (D2) days. (E) WT or Relb−/− (KO) cells transduced with EV or NFAT virus, differentiated in RANKL for 3 days and then infected with *S. aureus* and subjected to the gentamicin protection assay. Bars represent CFU of lysates at 18hpi. n=3 biological replicates. (A, B) ****p<0.0001 by two-way ANOVA with Tukey’s post-hoc test. (E)**p<0.01, ****p<0.0001 by one-way ANOVA with Tukey’s post-hoc test.

NFATc1 levels are significantly reduced with RelB deficiency, and retroviral overexpression of NFATc1 can rescue deficient OC formation that results from the loss of RelB [23]. Therefore, we next determined whether ectopic expression of NFATc1 could restore the ability of *S. aureus* to replicate in RelB-deficient cells. Consistent with a rescue in differentiation and NFATc1 restored to empty vector WT levels (Fig 4C, 4D), we found that microbial burden was also restored to empty vector WT levels in RelB knockout cells overexpressing NFATc1 under osteoclastogenic conditions (Fig 4E). These data highlight the importance of alternative NF-κB and NFATc1 signaling in mediating intracellular *S. aureus* replication.

### *S. aureus* within osteoclasts is not located exclusively in phagolysosomes

Macrophage subsets that kill bacteria after engulfment traffic them to phagolysosomes, a digestive hybrid organelle formed upon fusion of phagosomes with lysosomes [24]. To determine whether OCs fail to sequester *S. aureus* in phagolysosomes, we generated OCs on glass, infected with GFP+ bacteria, and stained the cells with the acidotrophic dye LysoTracker red at 18 hpi. Similar to the wide range in fluorescence intensity of infected D2 preOCs demonstrated by flow cytometry (Fig 2D), the number of bacteria in each mature OC was also variable (Fig 5A). Interestingly, in cells with a low bacterial load (Fig 5A, box 1), there is a high degree of colocalization between the bacteria and phagolysosomes (Fig 5B, 5C). Comparatively, cells harboring very large clusters of bacteria (Fig 5A, boxes 2 and 3) have more bacteria that do not appear to reside in acidified phagolysosomal compartments (Fig 5B), although the degree varies. The colocalization of the fluorescent signal peaks from the green GFP+ bacteria with the red LysoTracker stained phagolysosomes is represented by the Pearson’s colocalization correlation coefficients (Rr) generated from line scans taken within each cell (Fig 5C). The Pearson’s coefficients are inversely correlated to the amount of bacteria present within each of the three cells, indicating less phagolysosomal localization with higher bacterial loads. Thus, *S. aureus* within OCs seems to avoid killing by evading or escaping mature phagolysosomal compartments and the ensuing intraluminal digestion.

**Fig 5.**
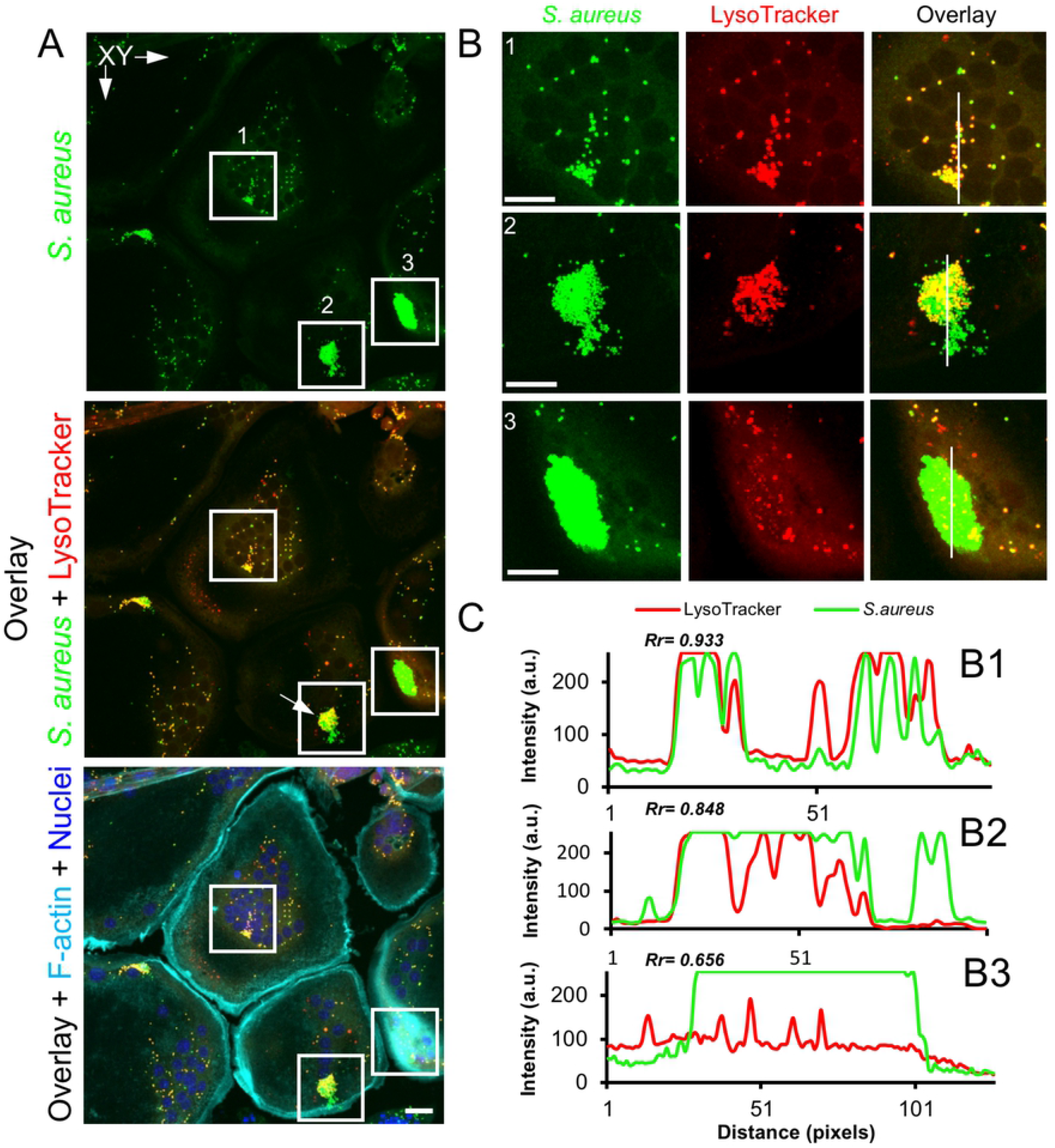
*S. aureus* in OCs is not exclusively in lysosomes. Confocal microscopy images of BMMs differentiated into OCs and then infected with GFP+ *S. aureus*. (A) Images show GFP+ *S. aureus* (green), lysosomes (LysoTracker, red), F-actin (turquoise), and nuclei (blue) within OCs. Enumerated inserts represent higher magnified images (B). (C) The cross-correlation analysis of the peaks of green *(S. aureus)* and red (lysotracker) fluorescence intensity from the line scans (B1-3, white lines) is represented with the corresponding Pearson’s coefficient (Rr).

## Discussion

OM is a common and debilitating infection, with associated osteolysis causing pain and pathologic fractures. To this point, most of the work examining OCs in the context of OM has focused on the stimulatory effect of *S. aureus* on OC activity. In contrast, we utilized a low multiplicity of infection (MOI) of 1 to 10 which may reveal host-pathogen interactions that are masked by much higher bacterial loads, as different *S. aureus* inocula have been shown to have differential effects on the immune response and infection progression [29]. Here, we have shown that OCs are a target of *S. aureus* infection, *in vivo* and *in vitro*, providing the bacteria a replicative niche. Unlike their progenitors, OCs are unable to confine internalized *S. aureus* to phagolysosomes, the likely cause of their failure to eliminate the bacteria. Thus, the direct interactions between OCs and *S. aureus* may play an important role in the progression of OM beyond bone loss, affecting the survival and proliferation of the pathogens.

The ability of *S. aureus* to proliferate after internalization was dependent on prior RANKL stimulation of the host cell for 2 days, the point at which cells become positive for TRAP and are considered to be committed to the OC lineage. Complete differentiation into multinucleated OCs, whether on plastic or bone, was associated with an even higher level of intracellular bacteria. The effect of RANKL was unique, as polarization of the BMMs toward M2 macrophages with IL-4, which causes reduced bacterial killing, did not allow intracellular replication. Since osteoclastogenesis depends on NFATc1, it is not surprising that this transcription factor was also required for the effect of RANKL on *S. aureus*. Interestingly, overexpression of NFATc1 alone, which is not sufficient to cause OC differentiation, did not promote bacterial proliferation without concomitant RANKL exposure. This suggests that additional signals from RANKL/RANK are required. The alternative NF-κB pathway is activated by RANKL and is upstream of NFATc1 and other factors important for OC function [23]. Deficiency of NIK, the apex kinase, results in a more severe impairment in OC differentiation relative to genetic disruption of RelB, the key transcriptional subunit [23], and likewise NIK ablation blunted *S. aureus* proliferation greater than RelB deficiency. Furthermore, restoration of NFATc1 expression in the RelB-deficient background to Wt levels also normalized both OC differentiation and bacterial loads, suggesting that in this context, NFATc1 activation is the primary mediator of bacterial handling. These findings are especially interesting in light of findings from others that elucidate the importance of RANKL-signaling in driving osteoclast-mediated inflammatory bone resorption [30].

Using a low MOI, we observed cellular heterogeneity within our cultures in both the D2 preOCs as well as in mature OCs. By flow cytometry, we were able to identify the small percentage of cells that actually became infected with *S. aureus*, and then follow the cultures over time. Since the fraction of infected cells did not change between 1.5 and 18 hpi, the increase in fluorescence represents bacteria replicating within previously infected cells, and not replication in the media and infection of new cells. We also found that amongst this infected population there were a subset of cells that allowed the replication of bacteria to a much greater extent than other cells, as reflected by the “tail” of high MFI cells only at 18 hpi in D2 preOCs. In fact, this minority population of cells could be responsible for the more than 2 log_10_ increase in bacteria represented in the MFI by flow cytometry and CFUs in the antibiotic protection assays. It is possible that differences in bacterial expansion within the preOCs at D2 are related to degree of differentiation, since these primary cell cultures are somewhat heterogeneous. However, a similar variability in bacterial load was seen by confocal microscopy in mature OCs.

As OCs are differentiated from BMMs, it is important to compare the consequences of bacterial internalization between the two cell types in order to understand how OCs fail where macrophages succeed. BMMs effectively destroy internalized bacteria via phagolysosome acidification. For *S. aureus* this is a process involving NLRP3 inflammasome activation and Caspase-1 cleavage that leads to NADPH oxidase 2 (NOX2) production of reactive oxygen species [24]. We discovered that most mature multinucleated OCs exhibited poor colocalization of *S. aureus* with phagolysosomes, and interestingly, the OCs with the highest intracellular bacterial loads had the lowest degree of *S. aureus* colocalization with phagolysosomes. Our initial ultrastructural investigation using transmission electron microscopy showed that dividing *S. aureus* can be found in membrane-bound compartments, but the bacterial load in the visualized cells was relatively low compared to most seen on confocal microscopy. Therefore, it is possible that bacteria replicate in the cytoplasm as well. Although in our experimental conditions lysis of OCs is uncommon before 18 hpi, it becomes frequent by 24 hpi. Future work will focus on tracing the intracellular fate of the bacteria within OCs over time, including defining the endosomal compartments involved, and whether they escape into the cytoplasm prior to cell death. Additional single cell approaches will be required to determine which host cell pathways determine the fate of intracellular *S. aureus*, and whether these are responsible for the observed differences between cells.

Ultimately, this work elucidates a new role for OCs in propagating infectious OM, at least in the context of *S. aureus*. It has previously been shown that osteoblasts release pro-osteoclastogenic cytokines, including RANKL, that recruit and activate OCs at the site of infection. While others have shown that *S. aureus* and its cellular components can promote osteoclastogenesis, OC activity, and pro-inflammatory cytokine release [17, 20, 31], this investigation focused on determining the infectability of OCs. We found that it is possible for OCs to become infected and provide a replicative niche for *S. aureus* to proliferate and evade immune destruction, and that this whole process is dependent on RANKL signaling. Although the influence of *S. aureus* on OC activity and cytokine production is undoubtedly important for the progression of OM lesions, we have extended the potential role for these cells to include bacterial expansion. If *S. aureus* promotes OC recruitment and formation and OCs can harbor bacteria from destruction, then these conditions can feed forward in a positive feedback loop. Thus, therapies aimed at modulating the ability of OCs to shelter bacteria might provide increased efficacy in curing these difficult-to-treat infections.

## Materials and Methods

### Reagents

Trypticase soy broth was procured from Fisher Scientific (Hampton, NH, USA). Fetal bovine serum and gentamicin (15750060) were purchased from Gibco-BRL (Grand Island, NY, USA). α-MEM, anti-α-actin (A2228) antibody, rosiglitazone, Polyinosinic-polycytidylic acid (Poly I:C), and lysostaphin (L7386) were all purchased from Sigma-Aldrich (St. Louis, MO, USA). Macrophage-colony stimulating factor (M-CSF), in the form of CMG 14-12 supernatant, and glutathione-S-transferase RANKL (GST-RANKL) were prepared as previously described [21]. Human M-CSF and Ficoll histopaque were purchased from Invitrogen (Carlsbad, CA, USA). Human CD14 magnetic beads were obtained from Miltenyi Biotec (Auburn, CA, USA). Anti-NFATc1 (7AG) antibody was obtained from Santa Cruz Biotechnology (Dallas, TX, USA) and anti-histone H3 (96C10) from Cell Signaling Technology (Beverly, MA, USA). Murine recombinant IL-4 and IFN-γ was purchased from Peprotech (Rocky Hill, NJ, USA).

### Mice

C57BL/6 male and female mice (8-10-week old) were purchased from The Jackson Laboratory (Bar Harbor, ME, USA). *Relb*^−/−^ and *Nik*^−/−^ mice and their littermate controls were generated by heterozygotic mating of *Relb*^+/−^ and *Nik*^+/−^, respectively, in a specific pathogen-free facility as previously described [22, 28]. TRAP promoter-tdTomato (TRAP^Red^) transgenic mice were generated by Dr. Ishii (Immunology Frontier Research Center, Osaka University, Osaka, Japan [27]), provided by Dr. J. Lorenzo, and maintained by heterozygotic mating in our specific pathogen-free facility. Conditional knockout *Nfatc1* (*Nfatc1 cKO*) mice were provided by Julia Charles (Harvard Medical School, Boston, MA) and generated by crossing *Nfatc1 fl/fl* mice with a transgenic line expressing the Cre recombinase from a type I interferon inducible promoter *(Mx1-Cre)* as previously described [32]. Activation of *Mx1-Cre* was achieved by intraperitoneal injection of 0.25 ml of 1 mg/ml poly I:C in PBS every other day. Three rounds of injections were given, followed by a three week waiting period to ensure optimal *Nfatc1* deletion. Littermates lacking the *Mx1-Cre* transgene were treated identically with poly I:C and served as NFATc1-sufficient (*Nfatc1fl/fl*) controls. For all experiments employing genetically modified mice, results were compared to cells from sex/age-matched littermate controls.

### Osteoclast culture

To generate osteoclasts (OCs) from enriched bone marrow macrophages (BMMs), bone marrow was harvested from the long bones of 10-12 week old mice and cells were cultured in α-MEM plus 10% FBS and a 1:10 dilution of CMG 14-12 cell supernatant (containing equivalent of 100 ng/ml of M-CSF) for 4 days to expand BMMs. Non-adherent cells were removed by several washes in PBS and adherent BMMs were detached with trypsin-EDTA, seeded into tissue-cultured treated plates and cultured in α-MEM plus 10% FBS containing a 1:50 dilution of CMG 14-12 cell supernatant (containing equivalent of 20 ng/ml of M-CSF) and GST-RANKL (60ng/mL) with media changes every day for indicated time periods.

For human osteoclastogenesis, peripheral blood mononuclear cells were obtained by density gradient centrifugation with Ficoll histopaque. Monocytes were isolated using anti-CD14 magnetic beads according to manufacturer directions. Human CD14+ cells were seeded into 6 well plates at a seeding density of 1 x 10^6^ cells per well. Monocyte-derived osteoclasts were generated by culturing CD14+ cells for 6 days in α-MEM plus 10% FBS containing 20 ng/ml of human M-CSF and GST-RANKL. Media was changed and cytokines were replenished every other day.

### Bacterial strains and growth conditions

All experiments were conducted with derivatives of *Staphylococcus aureus* USA300 clinical isolate LAC [33]. Bacterial strains were grown in trypticase soy broth (TSB) overnight at 37°C with shaking at 225 rpm, subcultured at a dilution of 1:100, grown to the mid-exponential phase (OD_600_ = 1.0) and centrifuged at 3000 rpm for 10 minutes. The pellets were washed and re-suspended with PBS to the desired concentration. To create a stable GFP+ strain of *S. aureus*, the region containing the *sarA* promoter driving sfGFP was first amplified out of pCM11 [34] using primers 5’-GTTGTT<u>TCTAGA</u>CTGATATTTTTGACTAAACCAAATG-3’ and 5’-GTTGTT<u>GAGCTC</u>TTAGTGGTGGTGGTG-3’ (restriction sites underlined). The resulting PCR amplification product was then ligated into the XbaI and SacI site of pJC1111 to create pNP1. pNP1 was then chromosomally integrated into the SaPI1 site of *S. aureus* as previously described [35]. The region encompassing the chromosomal SaPI1 integration was then transduced into *S. aureus* strain LAC (AH1263) [33] using phi80a. Integration of P*sarA*_sfGFP at the SaPI1 site was confirmed using primers JCO717 and 719 [35].

### Antibiotic protection assays

Infection of BMMs and OCs by *S. aureus* was quantified by determining the number of colony forming units (CFU) recovered from antibiotic treatment using a gentamicin protection assay as previously described [25, 26, 36]. Briefly, cells were seeded at 5 x 10^5^/well in 6-well plates and cultured in αMEM plus 10% FBS containing a 1:50 dilution of CMG 14-12 cell supernatant (containing equivalent of 20 ng/ml of M-CSF) in the absence or presence of GST-RANKL for the designated time of osteoclastic differentiation, as described above. To determine the level of intracellular survival, the cells were infected for 30 minutes at an MOI of 1:1 (final dose of 5 x 10^5^/well) at 37°C in 5% CO_2_, washed twice in PBS and cultured in media (α-MEM plus M-CSF +/− RANKL) containing antibiotic (gentamicin at final concentration of 0.3 mg/ml or lysostaphin at final concentration of 20ug/mL) for 1 hour to kill extracellular bacteria. Cells were washed twice in PBS to remove antibiotic and lysed in sterile, ice-cold ultra-pure H_2_0 for the 1.5 hour time point (1.5 hpi). For the 18 hour time point (18 hpi), culture media (α-MEM plus M-CSF +/− RANKL) was replaced after PBS washes and infection was continued to 18 hours post-infection prior to hypotonic lysis as described above. Lysates were 10-fold serially diluted, plated on TSB solidified with 1.5% agar (TSA), incubated overnight at 37°C and CFU enumerated. Controls for antibiotic killing of *S. aureus* were included in all experiments by plating supernatant on TSA and inspecting for colonies after overnight incubation at 37°C.

### Flow cytometric assays

For flow cytometry, cells were seeded at 5 x 10^5^/well in 6-well tissue culture treated plates, as described above. Cells were challenged with GFP-expressing *S. aureus* at an MOI of 1:1 for 30 minutes, and the extracellular bacteria was killed by the addition of gentamicin to the media for one hour. Cells were washed twice in PBS to remove antibiotic, and then media was replenished. The cells were then harvested at 2 hours (2 hpi) or 18 hours (18 hpi). Cells were detached with trypsin-EDTA, washed twice in PBS to remove non-adherent bacteria and extracellular fluorescence (reflecting attached but un-internalized bacteria) was quenched with 0.2% trypan blue as previously described [36] Cells were washed twice in PBS, fixed in 90% methanol for 30 minutes at 4°C and analyzed via flow cytometry (% FITC-positive cells and mean fluorescence intensity of FITC-positive population), using a BD LSR-II flow cytometer in 2mM EDTA, 2% FBS PBS. The data was analyzed using Flow Jo v 10.5.3.

### Confocal microscopy

Laser scanning confocal microscopy of mouse calvaria was performed to assess microbial uptake by osteoclasts *in vivo*. Prior to infection, RANKL (2 mg/kg body weight) was injected over the periosteum of calvaria of TRAP^Red^ reporter mice once per day for 5 days. Subsequently, 10^7^ GFP-expressing *S. aureus* were suspended in 100 μl PBS and subcutaneously injected over the periosteum of the calvaria. At 24 hours post-infection, calvariae were harvested, fixed overnight in 4% paraformaldehyde at room temperature and washed six times with PBS in 15 minute intervals. Calvariae were next decalcified in 14% free acid EDTA for 3 days under continuous agitation, infiltrated in 30% sucrose overnight at 4°C followed by embedding in OCT media and cut to 10 μm thick sections in a coronal orientation. Tissue sections were then mounted in Prolong Gold Antifade with DAPI and cured for 48 hours. For imaging of *in vivo* bacterial uptake, optical sectioning was performed by using Nikon A1RSi confocal microscope (Nikon Instruments Inc., NY, USA). Images were collected using an oil immersion 100x objective lens (CFI Plan Apo Lambda 100X Oil, Nikon Instruments Inc.) with 0.5 μm z-steps. For each mouse, 15-20 optical sections were captured in 8-10 different regions of interest from five separate tissue sections. Lasers at 405, 488, 561 nm were used to excite fluorescence from DAPI, GFP, and tdTomato reporter probes. Line-averaging and sequential scanning were employed to increase the signal-to-noise ratio and minimize spectral bleed-through, resulting in a frame acquisition time of 16 sec/frame. Background subtraction was carefully utilized to reduce the contribution of tissue autofluorescence. Images were processed in Nikon NIS-Elements (Nikon Instruments Inc.) and Fiji [37].

Murine bone marrow macrophage (BMM)-derived OCs differentiated on glass coverslips or devitalized bovine bone discs were incubated with GFP-expressing *S. aureus* (MOI=10) followed by gentamycin, as above, and cultured for a total of 18 hrs. To monitor for bacterial incorporation into phagolysosomes OCs were further incubated with the acidotrophic probe Lysotracker Red^®^ DND-99 (100 nm, Invitrogen) for 30 min prior to fixation. Cells were fixed in 4% paraformaldehyde (PFA) in PBS for 15 min at room temperature (RT), permeabilized with 0.1% Triton X-100 and stained with Alexa Fluor 647-conjugated phalloidin (1:500) and Hoechst 33258 dye (1:10,000) (Invitrogen) to visualize F-actin and nuclei, respectively. Samples were then mounted in ProLong^®^ Gold Antifade (Thermo Fisher Scientific Co.) and imaged using a Nikon A1RSi confocal microscope, equipped with a 10X (dry) lens and 60X (oil immersion) lens (Nikon Instruments Inc., NY, USA). Images were collected using the Nikon NIS-C Elements software. Cross-correlation analysis (Pearson’s, Rr) of colocalization between bacteria and phagolysosomes was performed using online ImageJ macros (NIH).

### Immunoblotting

Cells were washed twice on ice with cold PBS and lysed in RIPA buffer (20 mM Tris, pH 7.5, 150 mM NaCl, 1 mM EDTA, 1 mM EGTA, 1% Triton X-100, 2.5 mM sodium pyrophosphate, 1mM β-glycerophosphate, 1 mM Na3VO4, 1 mM NaF) containing Halt protease cocktail inhibitor (Thermo Scientific, Rockford, IL, USA). Following ten minutes incubation on ice, lysates were centrifuged at 16,000 x g to pellet cellular debris and protein concentration was determined by BCA quantification assay (Bio-Rad, Hercules, CA, USA). For each sample, thirty micrograms of total lysate was resolved by SDS-PAGE electrophoresis and semi-dry transferred to a PVDF membrane. Membranes were blocked for 1 hour in 5% milk in TBS containing 0.1% tween, probed with respective primary antibodies overnight at 4 °C with continuous agitation followed by 1 hour incubation at room temperature with secondary HRP-conjugated antibodies. Proteins were detected using WesternBright Quantum HRP substrate (Advansta, Menlo Park, CA, USA) and visualized with Genesnap using the Syngene Imaging System (Synoptics, Frederick, MD, USA).

### Retroviral transduction

NFATc1-pMX construct was transiently transfected into Plat-E packaging cells by calcium phosphate precipitation method as previously described [23, 38]. Viral supernatant was harvested 48 hours post-transfection and used to infect BMMs for 48 hours in the presence of M-CSF equivalent (20 ng/mL) and 4 μg/mL polybrene, followed by selection in blasticidin for 3 days before culture with 20 ng/mL M-CSF equivalent and GST-RANKL (30 ng/mL).

### Statistical Analysis

All data represented as the mean with standard deviation. Comparisons between groups of the CFUs from human OCs (Fig 2B) were analyzed by Student’s t-test. Comparisons between groups of CFUs from 18hpi OCs from different days of RANKL treatment (Fig 1D) or NFATc1 overexpression (Fig 4E) were analyzed by one-way ANOVA with Tukey’s multiple comparisons post-hoc test (GraphPad InStat). All other data analyzed by two-way ANOVA with Tukey’s multiple comparisons post-hoc test, and p<0.05 was taken as significant.

### Ethics Statement

All animal procedures were approved by Washington University Institutional Animal Care and Use Committees (IACUC Protocol Number 20170025), in compliance with the established federal and state policies outlined in the Animal Welfare Act (AWA) and enforced by the United States Department of Agriculture (USDA), Animal and Plant Health Inspection Service (APHIS), USDA Animal Care.

## Acknowledgments

This work was supported by National Institutes of Health grants R21 AR073507 and R01 AR070030 (to DJV), and by the Shriners’ Hospitals for Children grant 85117 (to DJV). JEC is supported by National Institute of Allergy and Infectious Diseases grants 1R01AI132560 and 5K08AI113107 and a Career Award for Medical Scientists from the Burroughs Wellcome Fund. NP is supported by NHMRC APP1143921. Histology was supported by the Washington University Resource Cores for Musculoskeletal Research (P30 AR074992). Confocal microscopy and data analysis were performed in part through the use of Washington University Center for Cellular Imaging (WUCCI) which is supported by Washington University School of Medicine, The Children’s Discovery Institute of Washington University and St. Louis Children’s Hospital (CDI-CORE-2015-505), the Foundation for Barnes-Jewish Hospital (3770) and the National Institute for Arthritis and Musculoskeletal Diseases (P30AR057235). We thank Linda Cox, Emily Goering, and Crystal Idleburg for technical support and laboratory contributions.

## Supporting Information

**S1 Fig. Osteoclasts show increased intracellular bacterial load 18 hours after *S. aureus* infection by lysostaphin protection assay.** Colony forming units (CFU) from lysates of OCs differentiated for 0, 2, or 3 days (D0, D2, D3, respectively) and subjected to the lysostaphin protection assay after infection with *S. aureus*. Lysates harvested at 18 hours post-infection. **p=0.0014, ****p<0.0001 by one-way ANOVA with Tukey’s post-hoc analysis. n=3 biological replicates.

**S2 Fig. OCs grown on bone chips *in vitro* allow *S. aureus* replication similar to OCs grown on plastic.** Colony forming units (CFU) from lysates of OCs grown on bone chips for 5 days and subjected to the gentamicin protection assay after infection with *S. aureus*, harvested at 1.5 or 18 hours post infection (hpi). **p<0.01 by t-test. n=3 biological replicates.

**S1 Table. Increased *S. aureus* colony formation is not the result of an increased extracellular bacterial load.** Colony forming units (CFU) from sampled media during the course of the gentamicin protection assay. Samples taken from the media of cells differentiated into OCs for 0 (D0) or 2 (D2) days at 12, 15, or 18 hours post infection (hpi). Colonies listed as total number of colonies formed (#) and the CFU (log) after 1000-fold dilution.

